# Identification of Cysteine-Reactive Early Mitotic Inhibitors

**DOI:** 10.64898/2026.10.13.758496

**Authors:** Xiaoxuan Zhang, Emma Buechler, Melissa Lopez, Mai Abdusamad, Cameron Taylor, Robert Damoiseaux, Jorge Z. Torres

## Abstract

Cancer remains a pervasive health problem for millions of Americans, and defining effective anticancer therapeutics remains a major obstacle to the long-term survival of patients. Recent advances in novel chemistries and chemical libraries have enabled researchers to discover and develop covalent inhibitors to treat various diseases, including cancer. This has spurred the generation of cysteine-targeting covalent compound libraries that have not been fully explored within the context of the cell cycle and cell division, which are attractive targets for inhibiting cancer proliferation. We report a novel, cell-based high-content chemical screening strategy for discovering early mitotic spindle assembly inhibitors. We coupled this strategy with a cysteine-reactive covalent compound library containing 3,520 molecules to identify novel early mitotic inhibitors. These inhibitors included a microtubule depolymerizer, a mitotic spindle positioning disruptor, and compounds affecting mitotic spindle structure and chromosome alignment. Two of these compounds triggered mitotic arrest and reduced cell viability at nanomolar concentrations. These inhibitors serve as chemical probes to investigate mechanisms of early cell division and to develop novel cancer therapeutics.

**Graphical Abstract:** 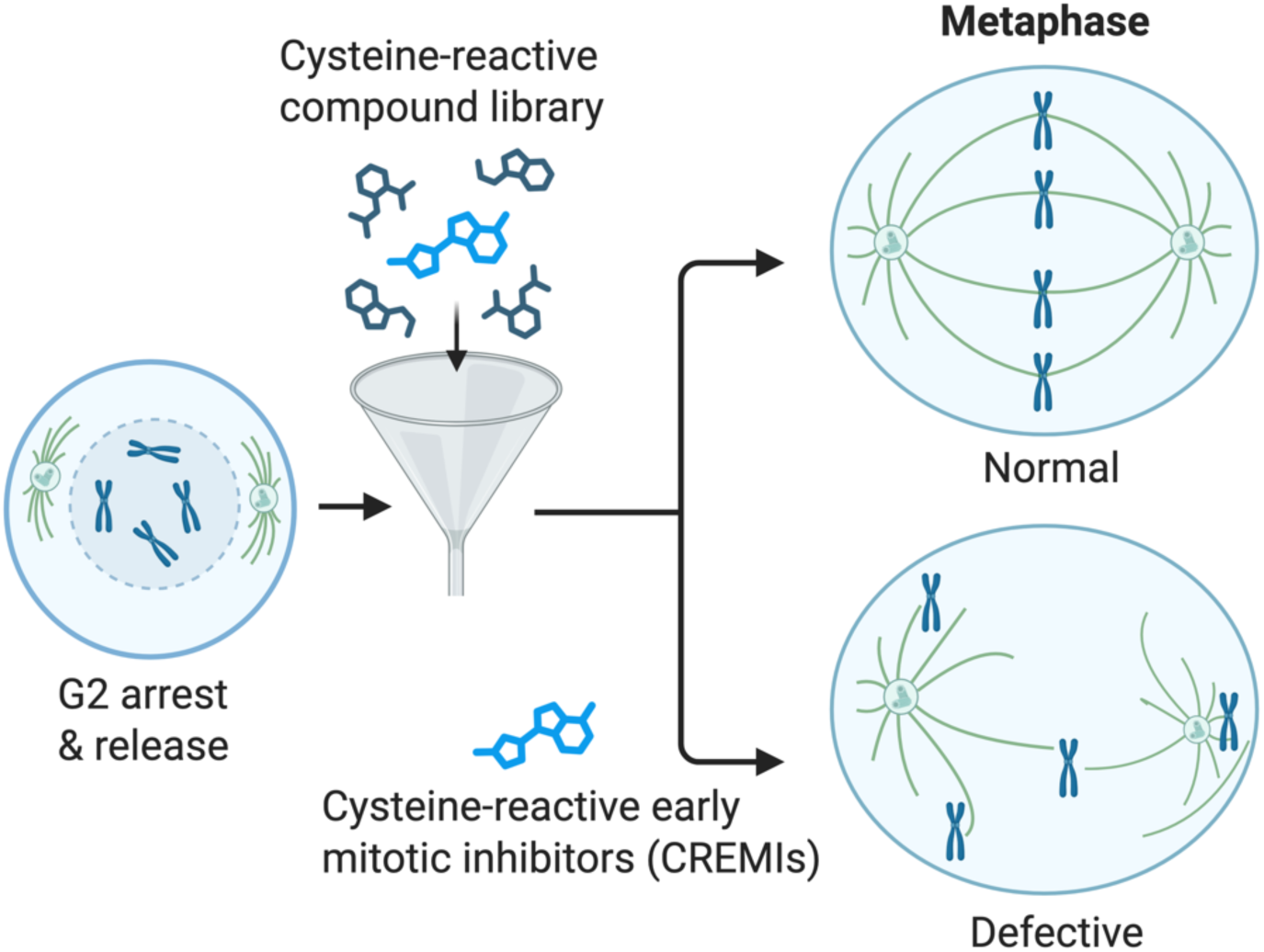

## INTRODUCTION

Over 2 million Americans will be diagnosed with cancer in 2026.^1^ Cancer is a proliferative disease, and pharmacological targeting of the cell division cycle that drives proliferation has been an intense area of study.^2, 3^ Although numerous chemical screens targeting the cell cycle have been conducted^4, 5^, few have focused specifically on the early, mid, or late subphases of cell division. Most screens have targeted the activity of specific cell division proteins through in vitro activity-based chemical screens and then used lead chemical inhibitors to study the effects of inhibiting these proteins during specific phases of cell division in cells. For example, Straight et al.^6^ screened for small-molecule inhibitors of nonmuscle myosin II and identified Blebbistatin, which they used to characterize myosin II function in cytokinesis, a late-stage process of cell division. Chemical inhibitors have advantages over traditional genetic approaches, such as RNAi and CRISPR-Cas9, in that they can inhibit protein function within seconds to minutes, compared to days with genetic approaches.^7, 8^ This has enabled researchers to utilize these chemical inhibitors to dissect cell division processes in a precise and temporal manner by coupling them to cell cycle synchronization approaches.

Traditional human cell cycle synchronization studies have used the washable DNA intercalating agent thymidine to arrest cells in S phase.^9^ Thereafter, cells are washed and released into the cell cycle and typically enter mitosis about eight hours later.^9^ More recently, the discovery of the reversible CDK1 inhibitor RO-3306 has allowed researchers to arrest cells at the G2/M transition, from which they can be released into early mitosis.^10^ Similarly, the proteasome inhibitor MG132 has been used to arrest cells with fully aligned chromosomes at the metaphase plate.^11, 12^ Coupling cell-synchronizing chemical treatments with specific enzyme inhibitors has also proven an effective approach for revealing previously intractable subphase-specific functions of key enzymes.^13^ For example, cell synchronization studies with the polo-like kinase 1 (PLK1) inhibitor BI2536, as performed in Brennan et al.^14^, showed that PLK1 was required for anaphase spindle elongation and proper cytokinesis.

It is estimated that the majority of the human proteome is “undruggable”, meaning it is difficult to target with traditional chemical inhibitors due to a lack of defined ligand-binding pockets.^15–17^ However, cysteine-directed covalent ligand drug discovery has emerged as a viable approach to target difficult-to-drug proteins, providing sustained inhibition and a longer residence time, as well as higher specificity due to the low abundance of accessible cysteines in the proteome.^16, 18^ This has sparked the recent development of several FDA-approved cysteine-directed covalent drugs for the treatment of various types of cancer, with many more being evaluated in clinical trials.^16^ For example, Sotorasib, a KRAS inhibitor, covalently binds to the cysteine of KRAS^G12C^ mutants, locking it in an inactive state.^16^

Currently, cell-based chemical screening campaigns for inhibitors targeting specific mitotic subphases are scarce, and there is a critical need to develop novel cancer therapeutics that leverage modern covalent chemical libraries to target proteins that have historically been difficult to drug. We report the development of an early cell division chemical screening approach that utilizes cell synchronizations, the latest developments in image-based artificial intelligence (AI) analysis software, and a recently developed cysteine-reactive covalent compound library to uncover novel early mitotic covalent inhibitors. Notably, we screened the kinome and cysteine-reactive libraries to identify 97 primary hit compounds, 88 of which reconfirmed on retesting, that inhibited or enhanced chromosome congression to the metaphase plate during early mitosis. Additionally, we performed a high-resolution multiparametric phenotypic analysis of cells treated with these inhibitors to classify their effects on mitosis, including microtubule depolymerization, defective chromosome congression, bypass of the MG132-induced metaphase arrest and transition into anaphase, and mitotic spindle positioning disruptors. Most of these inhibitors also arrested cancer cells in mitosis and reduced cell viability. These inhibitors serve as chemical leads for investigating the mechanisms of early spindle assembly and for developing anticancer therapeutics.

## RESULTS AND DISCUSSION

### Development of an Early Mitotic Inhibitor Screening Approach

Targeting early mitotic spindle assembly events is a viable strategy in the fight against proliferative diseases, such as cancer.^19^ The lack of precise chemical screening strategies targeting the early steps of mitosis prompted us to develop a mitotic subphase chemical screening approach to identify mitosis subphase-specific inhibitors (Figure 1 and Figure S1). As an overview, this approach utilized a cell cycle phase synchronization scheme where human HeLa cervical cancer cells plated in 384-well plates were arrested with RO-3306 for 16 h, washed and released into fresh media for 8 h, arrested with RO-3306 for 16 h, washed and released into MG132 for 2 h in the presence or absence of one compound/well from a chemical library (Figure 2A). Cells were then fixed and stained with the DNA dye Hoechst 33342, and their DNA morphology imaged with a high-content microscope (Figure 2B). Images were then analyzed with IN Carta artificial intelligence (AI) analysis software using two segmentation models. Model 1 was trained to identify the total number of cells (mitotic and non-mitotic), and Model 2 was trained to identify only cells arrested in metaphase with chromosomes aligned at the metaphase plate (Figure 2C). The percentage of cells arrested in metaphase was then quantified by dividing the total number of metaphase cells by the total number of cells. Statistical analyses were then conducted to identify compounds that altered the percentage of metaphase-arrested cells. Using the control microtubule stabilization drug taxol, which inhibits cells from forming a bipolar spindle and congression of chromosomes at the metaphase plate, this assay yielded a Z’ value of 0.49, indicating suitability for screening (Figure 2D).

**Figure 1.**
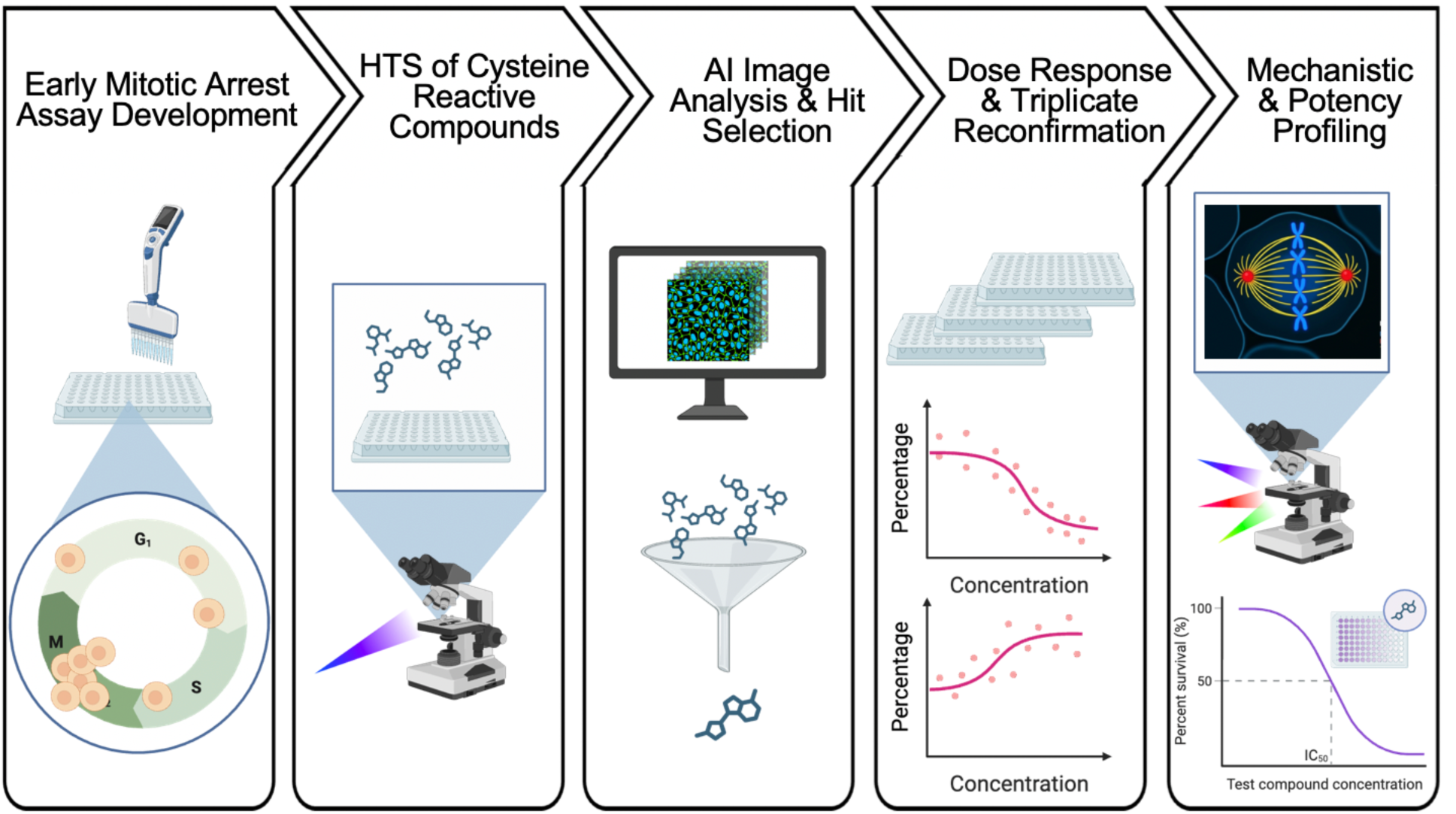
Overview of the identification and characterization of early mitotic covalent inhibitors. A double RO-3306 arrest and release cell synchronization strategy was coupled to a cell-based phenotypic high-throughput screening platform to screen a library of 3,520 cysteine-reactive small molecules. Two image analysis AI models were trained and applied to identify 16 primary hit compounds. A secondary validation screen in triplicate reconfirmed 12 compounds. A dose-response analysis was then used to determine the cell viability IC_50_ values for each compound. High-resolution immunofluorescence imaging was then used to quantify compound-induced mitotic phenotypes and investigate their mechanisms of action, followed by in vitro validation assays.

**Figure 2.**
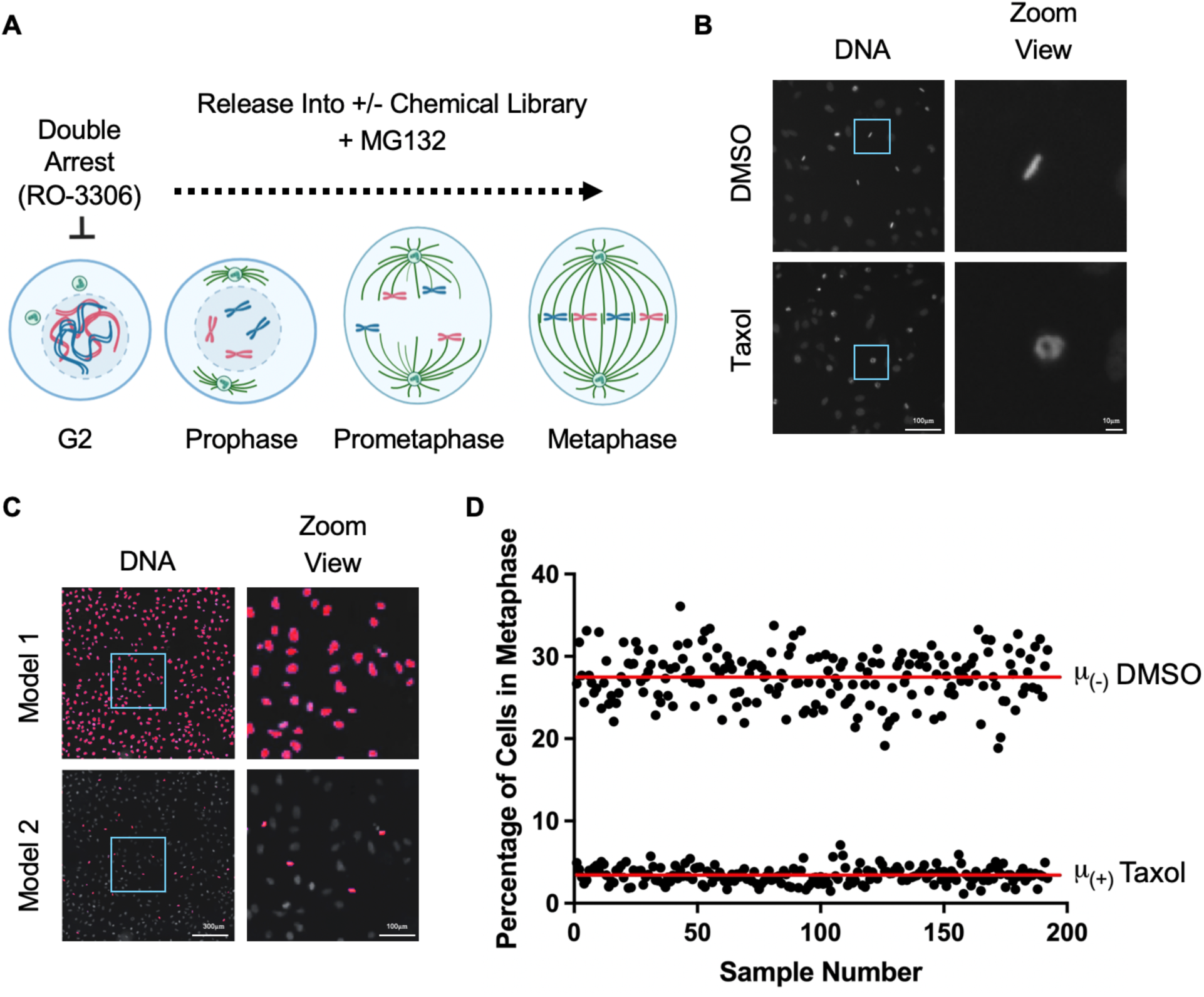
HeLa cells are plated onto 384-well plates, treated with the late G2 phase inhibitor RO-3306 for 16 h, released for 8 h, incubated with RO-3306 for 16 h, and released into mitosis for 2 h in the presence of MG132 and one drug/well from a compound library. (B) Images on the left show HeLa cells released into DMSO control or 100 nM taxol. Cells were fixed with 2% PFA, and the DNA was visualized with Hoechst 33342. Bar indicates 100 μm. Note that the addition of taxol induces a morphological deviation from the control metaphase-aligned chromosomes. The images on the right show a zoom view of the square area on the left image. Bar indicates 10 μm. (C) Two IN Carta deep learning AI models were trained to analyze images captured with an ImageXpress Confocal High-Content Imaging System. Model 1 recognizes and quantifies the total number of cells, while model 2 recognizes and quantifies cells with a metaphase morphology. The images on the right show a zoom view of the square area on the left image. Bars indicate 300 μm (left) or 100 μm (right). (D) HeLa cells in a 384-well plate were synchronized and released into mitosis, as in A, in the presence or absence of DMSO (192 wells) or 100 nM taxol (192 wells). For each well, the percentage of cells arrested in metaphase was calculated by dividing the number of cells in metaphase by the total number of cells. The solid red lines indicate the mean for all DMSO or taxol-treated samples.

### Early Mitotic Inhibitor Screening Approach Validation

Reversible post-translational modifications, in particular phosphorylation carried out by kinases, are critical for early spindle assembly, and numerous mitotic kinases have been identified to have a role in spindle assembly and chromosome congression to the metaphase plate.^20^ Therefore, we sought to expand the validation of this screening approach by screening through 2,589 compounds from the kinome library (MedChemExpress) at a final concentration of 500 nM. This analysis revealed 68 compounds with a sample Z score lower than −3 that inhibited metaphase chromosome alignment and 13 compounds with a sample Z score higher than +3 that enhanced it (Figure 3A and Table S1) with an overall hit rate of 3.1% (81/2,589). The inhibitors included compounds known to inhibit the activity of key mitotic kinases with important roles in cell division, such as Aurora kinases (NKIL-AMG-900, NKIL-AT9283, NKIL-VX-680, NKIL-SNS-314 Mesylate, MC-LY3295668, MC-BI-847325, MC-Ilorasertib, Hesperadin, and Aurora kinase inhibitor-8) and PLK1 (NKIL-Ro3280, Volasertib, BI 2536, MC-JNJ-10198409, MLN0905, HMN-176, and NKIL-GSK461364). Another example was the IGF-1R inhibitor MS-1504410 (Picropodophyllin) that inhibits centrosome separation and leads to monopolar spindle formation independent of the mitotic kinesin Kif11/Eg5.^21^ Also noteworthy was the identification of MC-XMD-17-51, an inhibitor of the microtubule-associated Doublecortin-like kinase 1 (DCLK1), which inhibits the proliferation and epithelial-to-mesenchymal transition (EMT) of lung cancer cell lines.^22^ Additionally, the transforming acidic coiled-coil 3 (TACC3) inhibitor MC-BO-264, which arrests cancer cell growth, especially cancers harboring the fusion oncogenic driver FGFR3-TACC3.^23^

**Figure 3.**
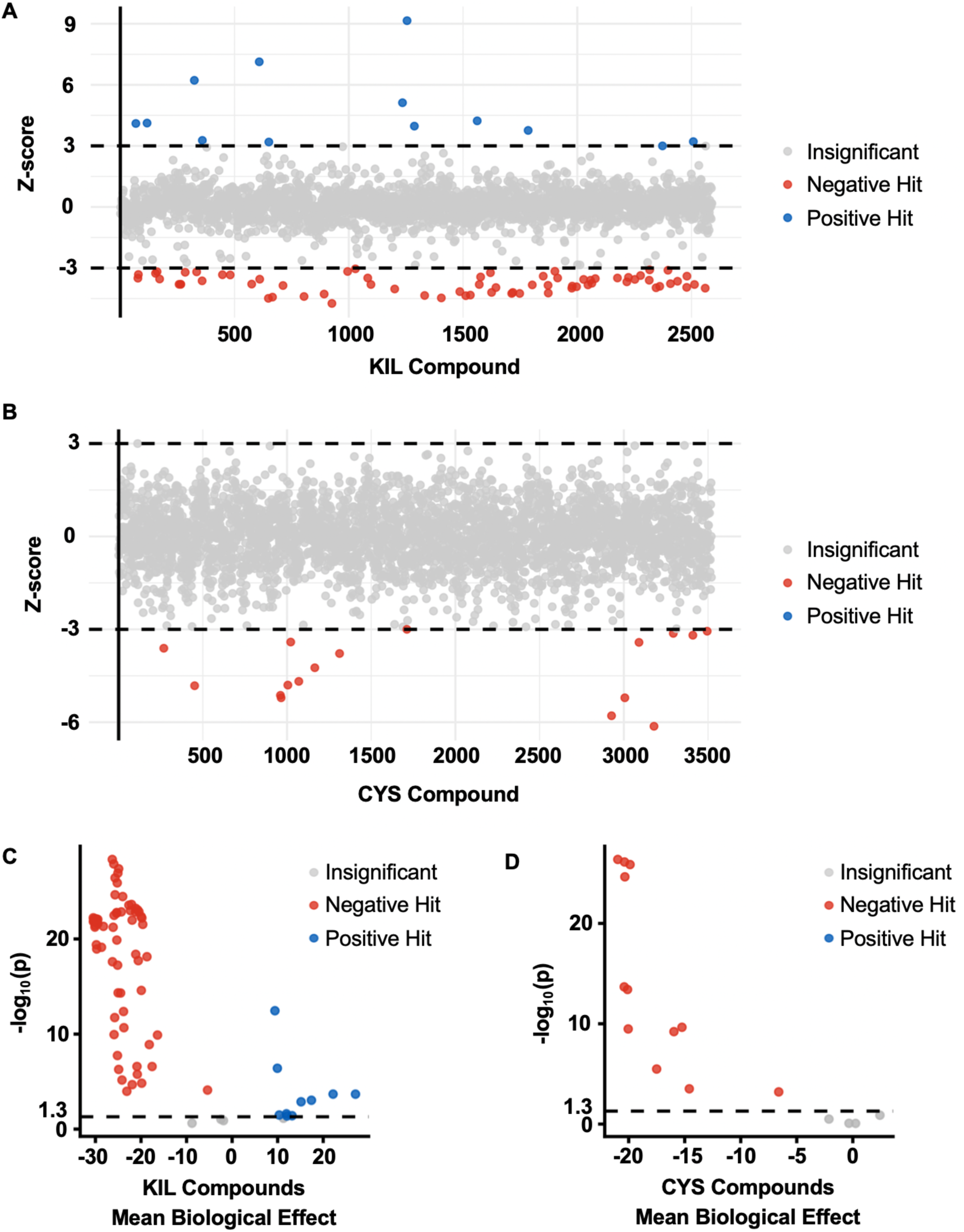
High-throughput screening of the kinome (KIL) and cysteine-reactive (CYS) chemical libraries. (A) The high-throughput screening workflow and automated image analysis pipeline described in Figure 2 were applied to screen the KIL library containing 2,589 small-molecule compounds. Sample Z-scores were calculated for each compound on a plate-by-plate basis. Negative hits (Z-score < −3) were compounds that decreased the percentage of metaphase-aligned cells. Positive hits (Z-score > 3) were defined as compounds that increased the percentage of metaphase-aligned cells. (B) The same workflow was applied to the CYS library containing 3,520 compounds using the same hit selection criteria, and only negative hits were identified. (C) Secondary reconfirmation of the primary KIL hits was performed by repeating the screen in triplicate, followed by a two-sided Student’s *t*-test comparing them with DMSO-treated controls on the same plate. A significance threshold of *p* < 0.05 was used. The mean biological effect was calculated as the difference in the percentage of metaphase-aligned cells between compound-treated and DMSO-treated wells. (D) Secondary reconfirmation of primary hits identified from the CYS library was performed using the same triplicate screening and statistical analysis described in (C).

The protein Tyrosine Kinase Wee1 is critical for establishing the G2/M checkpoint, and chemical inhibition of Wee1 leads to abrogation of the G2/M checkpoint.^24, 25^ Among the enhancers, we identified five Wee1 inhibitors (PD0166285, JND3229, WEE1-IN-3, PD166326, and Adavosertib), which are expected to enhance entry into mitosis upon release and to allow the accumulation of more metaphase cells within the assay timeframe (Figure 3A). Together, this screen indicated that our screening approach was robust, capable of identifying known inhibitors and enhancers of chromosome metaphase alignment, and capable of defining previously unappreciated compounds that modulated the ability of cells to progress properly through early mitosis.

### Discovery of Cysteine-reactive Early Mitotic Modulators

Despite the potential to develop cysteine-reactive compounds to treat proliferative diseases such as cancer, very few cysteine-reactive mitotic inhibitors have been identified. Therefore, we sought to expand the toolbox of cysteine-reactive inhibitors targeting early mitosis. To achieve this, we used our early-mitosis screening approach to test a library of 3,520 diverse compounds (Life Chemicals) featuring cysteine-reactive warheads, such as acrylamides, α,β-unsaturated ketones, and phenylsulfonate esters, at 2 μM. Our approach yielded 16 compounds with a sample Z score lower than −3 that inhibited metaphase chromosome alignment, but none that enhanced it (Figure 3B and Table S1) with an overall hit rate of 0.45% (16/3,520). The inhibitors included F0401-0081, F0482-0030, F0526-2186, F0526-2206, F0533-0686, F0777-2455, F0806-0881, F1190-0505, F3328-0012, F3382-3392, F3385-2487, F3385-4448, F5833-4735, F6122-0041, F6178-7509, F6468-0004, all of which had not been shown to affect mitosis (Table S1). Among these compounds, eight contained an acrylamide group, a classic Michael acceptor where the nucleophilic cysteine thiol attacks. Four compounds have a chalcone functional group, another type of Michael acceptor characterized by an α,β-unsaturated ketone conjugated to aromatic rings. Chalcones and their derivatives have been reported to interact with microtubules and inhibit tubulin polymerization, suggesting a potential mechanism underlying the anti-mitotic activity of these hit compounds.^26^ Two compounds have the exceptional leaving group sulfonate esters, enabling the covalent attachment of cysteine via an S_N_2 reaction. One compound contained a nitrile group and one contained a halogenated α,β-unsaturated ketone. Together, this screen identified novel cysteine-reactive compounds that inhibited cells from aligning chromosomes at the metaphase plate.

### Validation of Cysteine-reactive Early Mitotic Modulators

To further validate the inhibitors and enhancers of early spindle assembly identified in our kinome and cysteine-reactive screens, we retested all 97 hit compounds from the two screens (kinome 81, cysteine-reactive 16) that showed either a < −3 Z score or > +3 Z score change using the same assay at three different concentrations (kinome: 500 nM, 100 nM, and 16.6 nM; cysteine-reactive: 2 μM, 400 nM, and 100 nM) in triplicate for each condition (Figure 3C,D). A Student’s *t*-test was performed to compare the triplicate at each concentration with 16 DMSO-treated wells on the same plate. Overall, 91% (88/97) of the inhibitors and enhancers re-tested positive (*p*-value < 0.05) at one or more concentrations in our screen (kinome 76, cysteine-reactive 12) (Figure 3C,D and Table S2). The dose-response curves for metaphase-aligned cell number, total cell number, and percentage of metaphase-aligned cells were plotted. They revealed that compounds such as F1190-0505 from the cysteine-reactive library were likely toxic to cells, as a decrease in percentage of metaphase-aligned cells correlated with a decrease in total cell number (Figure 3C,D and Table S2). Additionally, compounds like F0526-2206 and F3328-0012 from the cysteine-reactive library showed the same significant effect on mitosis at both the lowest and highest concentrations tested (Table S2). Therefore, the optimal working concentration of each compound was determined as the lowest concentration required to induce robust early mitotic perturbation to minimize cytotoxicity and potential off-target effects (Table S2). Herein, we refer to <u>c</u>ysteine-<u>r</u>eactive <u>e</u>arly <u>m</u>itotic inhibitors as CREMIs.

### Microscopy Image-based Multiparametric Analysis of Cysteine-reactive Early Mitotic Inhibitors

Next, we sought to understand the mechanisms by which CREMIs were affecting early mitosis. HeLa cells were plated on coverslips, arrested in G2/M phase with 5 μM RO-3306, and then released into the cell cycle in the presence of one CREMI compound/well at the optimal concentration defined previously (Table S2), along with 10 μM MG132. Two hours later, cells were fixed and stained for DNA (Hoechst 33342), microtubules (rat anti-α-tubulin), spindle poles (mouse anti-γ-tubulin), and centrioles (rabbit anti-centrin-1), and imaged at 63X magnification by immunofluorescence (IF) microscopy. This multiparametric analysis of early mitosis revealed that all compounds induced a deviation from a normal mitosis. All CREMIs induced chromosome congression defects where chromosomes failed to align at the metaphase plate (Figure 4A,B and Table S3). Among them, five CREMIs also showed potential kinetochore-microtubule attachment defects, characterized by non-congressed chromosomes at spindle poles (Figure 4A,B and Table S3). Additionally, three CREMIs allowed cells to bypass the MG132-induced metaphase arrest and transition into anaphase (Figure 4A,C and Table S3). Together, these results indicated that hit CREMI compounds were likely inhibiting multiple targets in early spindle assembly.

**Figure 4.**
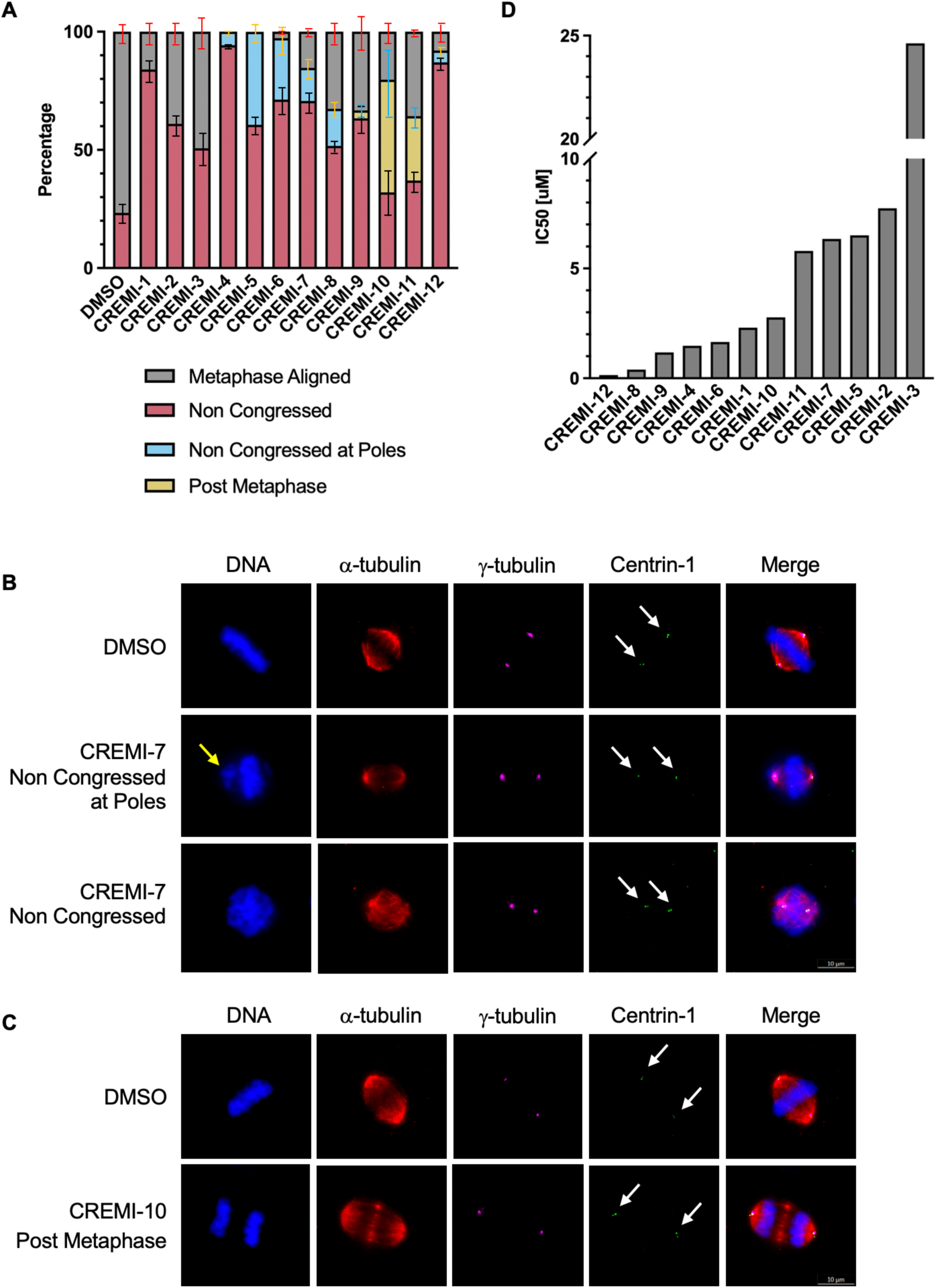
Quantitative analysis of the cellular effects of cysteine-reactive early mitotic inhibitor (CREMI) hit compounds. (A) High-resolution immunofluorescence imaging was used to characterize the identified CREMI hit compounds. Cells were fixed with 4% paraformaldehyde, permeabilized with 0.2% Triton X-100, and stained for DNA, α-tubulin, γ-tubulin, and centrin-1. For each compound, cells from three independent experiments were imaged, and 100 cells were quantified per coverslip. The bar plot shows the mean percentage of cells exhibiting each major mitotic phenotype across the three biological replicates for the 12 compounds. Most compounds predominantly induced non-congressed chromosomes, while a subset also caused chromosomes to scatter at the spindle poles or bypass metaphase and arrest in anaphase or telophase. The y-axis indicates the percentage of cells with the indicated cell division defects for the compounds tested on the x-axis. (B-C) Representative images of the three major phenotypes. Arrows indicate the position of the centrioles (centrin-1 staining). Bars indicate 10 μm. (D) HeLa cells were treated with increasing concentrations of each CREMI hit compound for 72 h, and cell viability was assessed using the CellTiter-Glo assay. IC_50_ values were determined by nonlinear regression analysis using a four-parameter logistic model. The y-axis indicates cell viability IC_50_ [μM] for the compounds indicated on the x-axis.

### Potency Analysis of Cysteine-reactive Early Mitotic Inhibitors

Next, we determined the potency of CREMI compounds for inducing cell death. HeLa cells were treated with a fourteen-step series of two-fold dilutions (100 μM to 12.2 nM) of each CREMI for 72 h, and the cell viability half-maximal inhibitory concentration (IC_50_) was determined using the CellTiter-Glo luminescent cell viability assay. This approach showed that most CREMIs were potent with cell viability IC_50s_ < 7 μM (Figure 4D and Table S4). CREMI-12 and CREMI-8 were particularly potent with IC_50s_ of 134 nM and 398 nM, respectively.

### CREMI-12 Perturbs Mitotic Spindle Positioning

In vitro, most proliferating cell lines undergo symmetric cell division to produce two identical cells with the same size and genetic material.^27^ Interestingly, we noticed that unlike DMSO-treated control cells that displayed a centrally positioned spindle, cells treated with 100 nM of CREMI-12 positioned their spindle towards the cell periphery of one of their spindle poles (Figure 5A). Quantitative analysis showed that 94.33 ± 3.06% (mean ± SD, n=3) of mitotic spindles were centrally positioned, with the spindle center within 25% of the cell centroid, in DMSO-treated cells, while only 19.33 ± 6.66% (mean ± SD, n=3) of mitotic spindles were centrally positioned in CREMI-12-treated cells (Figure 5B, Table S3). This analysis suggests that CREMI-12 likely targets a component critical for mitotic spindle positioning.

**Figure 5.**
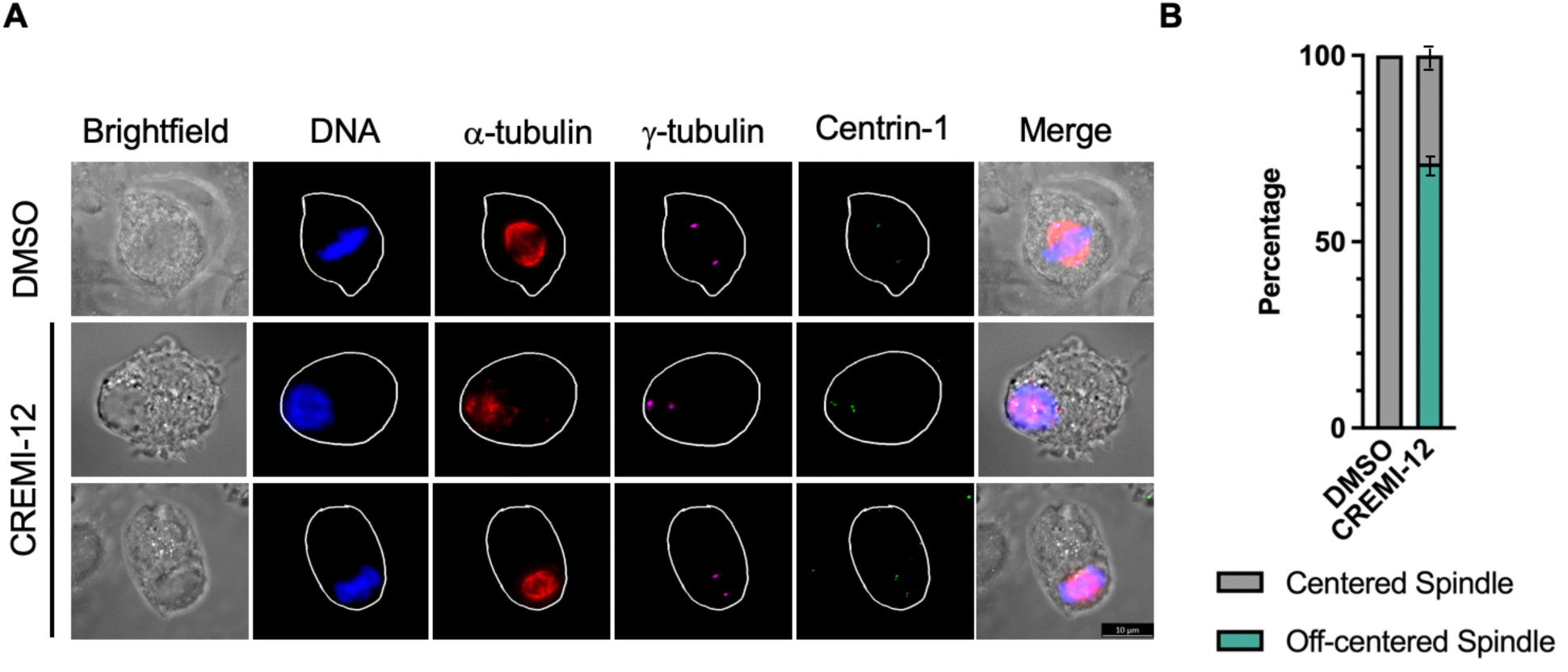
CREMI-12 perturbs the central positioning of the mitotic spindle. (A) Representative immunofluorescence images of HeLa cells treated with DMSO or 100 nM CREMI-12 and fixed and stained for DNA, α-tubulin, γ-tubulin, and centrin-1. Cell outlines were traced from the brightfield channel and overlaid on all other channels. Note that treatment with CREMI-12 led to the mitotic spindle shifting away from the center of the cell to the periphery. Bar indicates 10 μm. (B) Cells from three independent experiments were imaged, and 100 cells were quantified per experiment. Bar plot displays the average percentages of the major phenotypes across the three replicates (y-axis) for each treatment (x-axis).

### CREMI-1 Inhibits Microtubule Polymerization

Microtubules are a primary target for inhibiting cell division with small molecules^28^; however, few cysteine-reactive microtubule-targeting compounds have been identified. In our analysis, HeLa cells treated with 2 μM CREMI-1 displayed a microtubule-depolymerization phenotype (Figure 6A). Therefore, we tested whether CREMI-1 could inhibit tubulin polymerization in an in vitro tubulin polymerization assay. Briefly, we performed a fluorescence-based tubulin polymerization assay using control DMSO, 20 µM CREMI-1, 3 µM taxol (tubulin-polymerizer), or 3 µM nocodazole (tubulin-depolymerizer). We captured fluorescence measurements every minute for 90 minutes (Figure 6B). While taxol induced rapid tubulin polymerization, both CREMI-1 and nocodazole inhibited tubulin polymerization (Figure 6B). These results indicated that CREMI-1 targeted tubulin and inhibited its polymerization.

**Figure 6.**
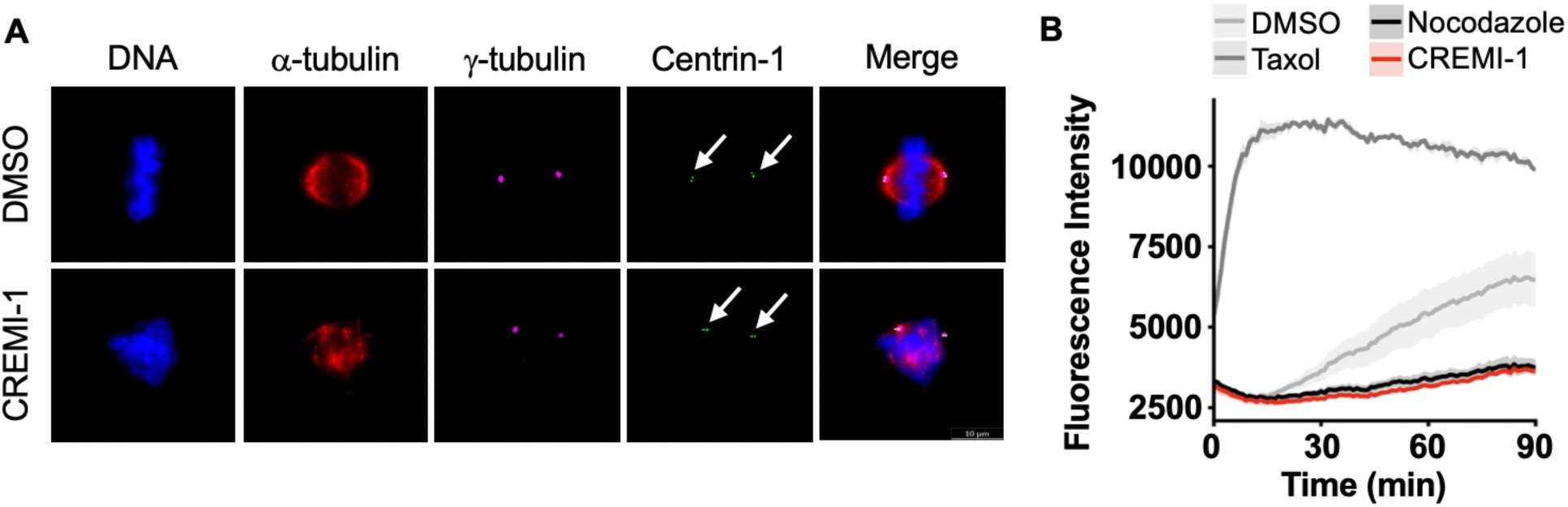
CREMI-1 inhibits microtubule polymerization. (A) Representative immunofluorescence images of HeLa cells treated with DMSO or 2 μM CREMI-1 and fixed and stained for DNA, α-tubulin, γ-tubulin, and centrin-1. Arrows indicate the position of the centrioles (centrin-1 staining). Bar indicates 10 μm. (B) *In vitro* microtubule polymerization reactions were carried out in the presence of DMSO, 3 μM taxol, 3 μM nocodazole, or 20 μM CREMI-1 at 37°C for 90 minutes, with fluorescence intensity measurements captured every minute. The experiment was repeated in triplicate, and the mean with standard deviation ribbon at each time point was plotted. The graph displays the mean fluorescence intensity (y-axis) over time (x-axis) for the indicated treatments.

## CONCLUSIONS

Cancer is a public health crisis and a leading cause of death in America.^29^ Cysteine-reactive compounds offer novel vantage points for targeting the molecular machinery responsible for cancer cell proliferation.^15–18^ Here, we coupled cell synchronization studies, diverse cysteine-reactive compounds, high-content image-based microscopy, and automated AI image analysis to define novel cysteine-reactive early mitotic inhibitors (CREMIs). These CREMIs included novel compounds that affected mitotic spindle assembly, positioning, and architecture, leading to mitotic arrest and subsequent cell death. Although the compound library was assembled around cysteine-reactive warheads, covalent engagement remains to be established. Future studies could focus on target deconvolution for 11 of 12 orphan CREMIs. Nonetheless, these CREMIs represent potential covalent leads to interrogate early spindle-assembly mechanisms and to develop new classes of therapeutics to treat cancer.

## MATERIALS AND METHODS

### Cell Culture

HeLa cells, purchased from and authenticated by the ATCC, were cultured in phenol red-free DMEM/F-12 1:1 and 10% fetal bovine serum at 37°C with 5% CO_2_. Media were purchased from Gibco. See Table S5 for a list of key reagents and resources used in this study.

### Screening Conditions

The screening assay was carried out in 384-well plates. A MultiFlo dispenser (BioTek) was used to add 10 μL/well of media to column 1 as a no-treatment control, 10 μL/well of RO-3306 (Selleckchem) solution to columns 2-24 (final 5 μM), and 15 μL/well of HeLa cell suspension to all wells (final 1,875 cells/well). Plates were incubated at 37°C in 5% CO2 for 16 h. An EL406 plate washer (BioTek) was used to wash the plates twice with 50 μL/well of media; 30 μL/well remained after the final wash. The plates were incubated at 37°C in 5% CO2 for 8 h. The MultiFlo was then used to add 30 μL/well of media to column 1 and 30 μL/well of RO-3306 to columns 2-24 (final 5 μM). The plates were incubated at 37°C in 5% CO2 for 16 h. The EL406 was used to wash the plates twice with 50 μL/well of media, and 30 μL/well was left after the final wash. The Beckman Coulter Echo 555 was then used to transfer 60 nL/well DMSO (Invitrogen) to columns 2 and 23 (final 0.2%), 60 nL/well taxol (Tocris) to column 24 (final 100 nM), 30 nL/well MG132 (Selleckchem) to columns 2-24 (final 10 μM), and each compound from the indicated compound library (45 nL/well kinome, final 500 nM; 60 nL/well cysteine-reactive, final 2 μM) to columns 3-22. The plates were incubated at 37°C in 5% CO2 for 2 h. The MultiFlo was used to add 30 μL/well of fix and stain solution to all wells (final 4 μg/mL Hoechst 33342 (Thermo Fisher Scientific) and 2% PFA (Electron Microscopy Sciences)). The plates were sealed with aluminum foil to shield from light and incubated at room temperature for 1 h prior to imaging.

### High-Content Imaging

Screen plates were imaged with an ImageXpress Confocal High-Content Imaging System (Molecular Devices) running MetaXpress 6.7. Acquisition used the wide-field configuration and a 10×0.45 NA air objective, with 4 sites imaged per well. The Hoechst 33342 DNA stain was visualized at 377 nm excitation and 447 nm emission. Camera binning was set to none, yielding 0.662 × 0.662 µm/pixel. Laser power and exposure were set on DMSO and taxol control wells such that no channel contained saturated pixels, and all acquisition settings were held constant across every plate in the screen.

### AI Model Training

We developed two SINAP-based segmentation models in IN Carta v2.7. AI image analysis software (Molecular Devices). The training dataset comprised more than 10,000 cells from 1,538 images acquired on the ImageXpress Confocal across six independent experiments. SINAP models are U-Net-style encoder–decoder convolutional networks with a ResNet encoder; the architecture is proprietary, and the vendor provided this description. We generated Model 1 by fine-tuning the Nuclei.a base model and used it to identify and quantify the total cell population, including both mitotic and non-mitotic cells. We set the minimum area to 50 µm^2^ to exclude dead cells and debris. Model 2 was custom-trained to identify cells arrested in metaphase with chromosomes aligned at the metaphase plate. We manually annotated images to label all metaphase-aligned cells, and incorporated these annotations into the SINAP training dataset for machine-learning segmentation. We generated six model iterations and used the final iteration for all reported analyses. An investigator manually assessed model performance on a set of 4 images (1,000 cells) drawn from DMSO- and taxol-treated control wells that were excluded from all rounds of model training. Model 1 achieved 98.72% precision, 97.47% recall, and 98.09% F1 score, while model 2 achieved 93.1% precision, 95.29% recall, and 94.19% F1 score. Both models were consistent and reliable across DMSO, taxol, and drug conditions. PhenoGlyph phenotypic classification was evaluated as an additional analysis layer following SINAP segmentation but did not improve classification performance; SINAP segmentation alone was therefore used for all reported analyses.

### Image Analysis

Images were imported into IN Carta, and both models were applied sequentially. Model 1 quantified the total number of cells, whereas Model 2 quantified the number of metaphase-aligned cells. The results were exported to Microsoft Excel, and the percentage of metaphase-aligned cells for each well was calculated by dividing the number of metaphase-aligned cells identified by Model 2 by the total cell count identified by Model 1.

### Assay Performance

To evaluate assay performance, a 384-well plate was prepared in which 192 wells received the screening condition containing 0.2% DMSO (negative control) and the remaining 192 wells received the screening condition containing 100 nM taxol (positive control). The quality of the assay was assessed by calculating the Z′ factor using the following equation:

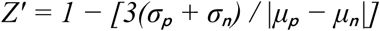

where σ represents the standard deviation and µ represents the mean of the positive (p) and negative (n) controls. The screening assay achieved a Z′ factor of 0.49, indicating usability for screening purposes.

### Primary Screen Hit Selection

The kinome library (KIL, 2589 compounds) and the cysteine-reactive library (CYS, 3520 compounds) were screened at 500 nM and 2 µM, respectively, in single wells. Each 384-well plate included 16 wells with DMSO and 16 wells with 100 nM taxol as in-plate controls. For each compound treatment, we calculated the percentage of metaphase-aligned cells. Sample Z scores were then calculated on a plate-by-plate basis according to the following equation:

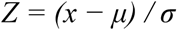

where x is the percentage of metaphase-aligned cells for an individual compound-treated well, µ is the mean percentage of metaphase-aligned cells across all compound-treated wells on the same plate, and σ is the corresponding standard deviation. Compounds with Z scores greater than 3 or less than −3 were considered primary hits. Data analysis was performed using Collaborative Drug Discovery (CDD Vault), and the results were exported as a Microsoft Excel file.

### Hit Triplicate Reconfirmation

Candidate hits identified in the primary screen were re-evaluated in triplicate using three concentrations (kinome library: 500 nM, 100 nM, 16.6 nM; cysteine-reactive library: 2 µM, 400 nM, 100 nM). For each compound, statistical significance was assessed using a two-sided Student’s *t*-test assuming equal variance, comparing the three replicate wells at each concentration with DMSO control wells from the same plate. Dose-response curves were generated using CDD Vault, and the resulting data were exported to Microsoft Excel for further analysis. Based on the dose-response analysis and microscopic evaluation of cellular morphology, we defined the optimal working concentration for each compound as the lowest concentration required for a significant *t*-test with a *p*-value < 0.05.

### Cell Viability Assays

HeLa cells were plated at 7500 cells/well in 96-well plates, and compounds were added 24 h after plating; each condition was tested in triplicate using a 14-point titration (12.2 nM to 100 µM). After 72 h of drug treatment, cell viability for each compound was calculated as described previously^4^ using the CellTiter-Glo assay (Promega). IC_50_ values were determined by nonlinear regression analysis using a four-parameter logistic model in GraphPad Prism.

### Tubulin Polymerization Assay

Tubulin polymerization assays were performed as described previously^30^ using the Tubulin Polymerization Assay Kit (Cytoskeleton). Reactions were carried out in the presence of DMSO, 3 µM taxol, 3 µM nocodazole, or 20 µM CREMI-1. Microplates were incubated at 37 °C for 90 min, and total fluorescent intensity was measured at 1-min intervals using a FlexStation 3 Multi-Mode Microplate Reader (Molecular Devices). Each condition was repeated in three independent experiments. The mean ± standard deviation at each time point was calculated and plotted using R.

### Fixed-cell Immunofluorescence

HeLa cells were fixed and imaged by immunofluorescence microscopy as described previously^31^, except that images were captured with a Leica MICA microscope (Leica Microsystems, 63x/1.40 NA oil objective, Leica Application Suite X software) and exported as .tiff files. For quantifying off-center spindles, spindles were considered off-center if the spindle center was >25% from the cell centroid.

### Antibodies

α-tubulin (Bio-Rad cat#MCA77G); γ-tubulin (Abcam cat#ab11316); centrin-1 (Proteintech cat#12794-1-AP); FITC, Cy3, and Cy5 conjugated secondary antibodies were from Jackson Immuno Research.

## ASSOCIATED CONTENT

### Supporting Information

The Supporting Information is available free of charge on the publications website. Supporting Information Figure S1 and Tables S1, S2, S3, S4, and S5.

### Data Availability

The data underlying this study are available in the published article and its Supporting Information. Raw microscopy images are available from the corresponding author upon reasonable request.

## AUTHOR INFORMATION

### Corresponding Authors

* Jorge Torres, Department of Chemistry and Biochemistry, UCLA, Los Angeles, CA 90095. Phone: 310-206-2092.

* Robert Damoiseaux, Department of Molecular and Medical Pharmacology, UCLA, Los Angeles, CA 90095. Phone: 310-794-1974.

### Author Contributions

X.Z., E.B., M.L., M.A., C.T., R.D., and J.Z.T. contributed to project design, chemical screening, data analysis, and manuscript preparation. X.Z. and E.B. performed cell and molecular biology experiments.

### Notes

The authors declare no competing financial interests.

## ACKNOWLEDGEMENTS

This work was supported by NIH-NIGMS R35GM139539 (J.Z.T) and NIH-NIGMS T32GM145388 (E.B. and M.A.) grants. Work conducted in the UCLA Molecular Screening Shared Resource was supported by NIH-NCI P30CA016042. The content is the responsibility of the authors and does not represent the official views of the National Institutes of Health.

