## Supplemental Material for "Identification of Cysteine-Reactive Early Mitotic Inhibitors"

### Table of Contents

Figure S1. Overview of the early mitotic inhibitor screening approach used for identification of cysteine-reactive early mitotic inhibitors (CREMIs)…………………………………………..…..S3

Table S1. Primary chemical genomic screens hits………..………………...………….………......S4

Table S2. Screen hit triplicate revalidation data and CREMI compound purity information…...S4

Table S3. CREMI compound treatment phenotype quantification……………….…………….S4

Table S4. CREMI compound cell viability $IC_{50s}$……………………………………………….…….S4

Table S5. List of reagents and resources used in this study……………………………………..S5

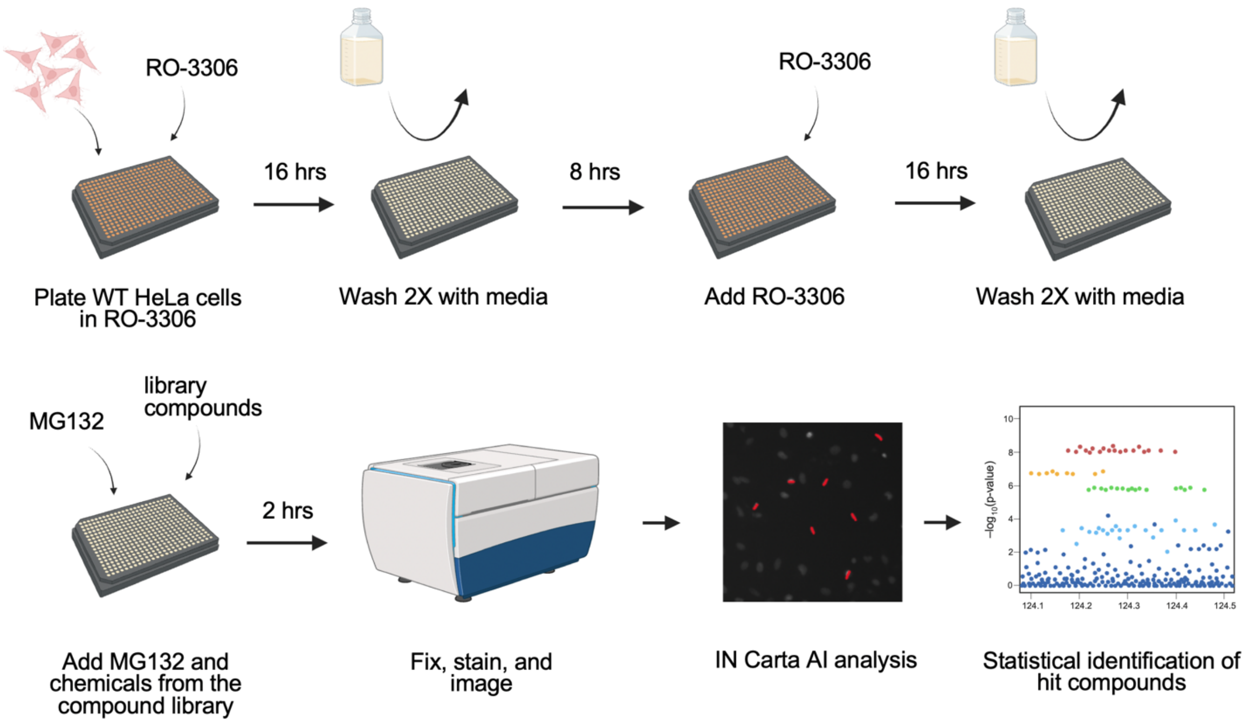


**Figure S1.** Overview of the early mitotic inhibitor screening approach used for identification of cysteine-reactive early mitotic inhibitors (CREMIs).

**Table S1.** Data summary and statistical significance analysis of chemical genomic screening data used to identify hit early mitotic inhibitors.

**Table S2.** Data summary and statistical significance analysis of hit early mitotic inhibitors revalidated in triplicate along with purity information for the 12 cysteine-reactive early mitotic inhibitors (CREMIs).

**Table S3.** Data summary and statistical significance analysis of cell division defects induced by CREMI compounds.

**Table S4.** Data summary and statistical significance analysis of cell viability IC50s for each of the CREMI compounds.

**Table S5.** List of reagents and resources used in this study.

**Table S5.** List of reagents and resources used in this study.

| REAGENT or RESOURCE | SOURCE | IDENTIFIER |
|---|---|---|
| Chemicals | | |
| RO3306 | Selleckchem | Cat# S7747<br>CAS: 872573-93-8 |
| DMSO | Invitrogen | Cat# D12345<br>CAS: 67-68-5 |
| Taxol | Tocris | Cat# 1097<br>CAS: 33069-62-4 |
| MG132 | Selleckchem | Cat# S2619<br>CAS: 1211877-36-9 |
| Hoechst 33342 | Thermo Fisher Scientific | Cat# H1399<br>CAS: 23491-52-3 |
| Paraformaldehyde | Electron Microscopy Sciences | Cat# 15710-S<br>CAS: 30525-89-4 |
| ProLong Gold Antifade Mountant | Thermo Fisher Scientific | Cat# P36934 |
| CellTiter-Glo | Promega | Cat# G9241 |
| Assay Kits | | |
| Tubulin Polymerization Assay Kit | Cytoskeleton, Inc. | Cat# BK011P |
| Antibodies | | |
| Tubulin Alpha antibody | Bio-Rad | Cat# MCA77G<br>RRID: AB_325003 |
| Gamma Tubulin antibody [GTU-88] | Abcam | Cat# ab11316<br>RRID: AB_297920 |
| Centrin 1 Polyclonal antibody | Proteintech | Cat# 12794-1-AP<br>RRID: AB_2077371 |
| Donkey polyclonal anti-Rat IgG (H+L), Cy3 AffiniPure | Jackson ImmunoResearch Labs | Cat# 712-165-153<br>RRID: AB_2340667 |
| Donkey polyclonal anti-Mouse IgG (H+L), Cy5 AffiniPure | Jackson ImmunoResearch Labs | Cat# 715-175-151<br>RRID: AB_2340820 |
| Donkey polyclonal anti-Rabbit IgG (H+L), Fluorescein (FITC) AffiniPure | Jackson ImmunoResearch Labs | Cat# 711-095-152<br>RRID: AB_2315776 |
| Cell Lines and Media | | |
| HeLa cells | ATCC | Cat# CCL-2<br>RRID: CVCL_0030 |
| DME/F-12 1:1 (1X) | Cytiva | Cat# SH30023.02 |
| DMEM/F12, no phenol red | Gibco | Cat# 21041025 |
| 10% FBS | Gibco | Cat# 16000044 |
| 0.25% Trypsin-EDTA (1X) | Gibco | Cat# 25200-056 |
| Software and Algorithms | | |
| MultiFlo FX Multimode Dispenser | BioTek | RRID:SCR_019746 |
| EL406 Washer Dispenser | BioTek | RRID:SCR_019735 |
| Echo 555 Liquid Handler | Beckman Coulter | https://www.beckman.com/liquid-handlers/echo-for-regulated-labs |
| ImageXpress Confocal High-Content Imaging System | Molecular Devices | RRID:SCR_020294 |
| FlexStation 3 Multi-Mode Microplate Reader | Molecular Devices | RRID:SCR_025281 |

| | | |
|---|---|---|
| IN Carta Image Analysis Software | Molecular Devices | https://www.moleculardevices.com/products/cellular-imaging-systems/high-content-analysis/in-carta-image-analysis-software |
| Collaborative Drug Discovery Vault | Collaborative Drug Discovery | https://www.collaborativedrug.com |
| BioRender | BioRender | RRID: SCR_018361 |
| Leica Mica Widefield Live Cell Microhub Automated Microscope | Leica Microsystems | https://www.leica-microsystems.com/products/confocal-microscopes/p/mica-widefield-live-cell/ |
| Leica Application Suite (LAS) X | Leica Microsystems | RRID:SCR_013673 |
| Tecan Infinite M1000 Microplate Reader | Tecan | RRID:SCR_025732 |
| RStudio 2024.09.1+394 | The R Project for Statistical Computing | RRID:SCR_001905 |
| GraphPad Prism 11 | GraphPad | RRID: SCR_002798 |
